# Underground deserts below fertility islands? – Woody species desiccate lower soil layers in sandy drylands

**DOI:** 10.1101/2020.01.20.912220

**Authors:** Csaba Tolgyesi, Peter Torok, Alida Anna Habenczyus, Zoltan Batory, Valko Orsolya, Balazs Deak, Bela Tothmeresz, Laszlo Erdos, Andras Kelemen

## Abstract

Woody plants in water-limited ecosystems affect their environment on multiple scales: locally, natural stands can create islands of fertility for herb layer communities compared to open habitats, but afforestation has been shown to negatively affect regional water balance and productivity. Despite these contrasting observations, no coherent multiscale framework has been developed for the environmental effects of woody plants in water-limited ecosystems. To link local and regional effects of woody species in a spatially explicit model, we simultaneously measured site conditions (microclimate, nutrient availability and topsoil moisture) and conditions of regional relevance (deeper soil moisture), in forests with different canopy types (long, intermediate and short annual lifetime) and adjacent grasslands in sandy drylands. All types of forests ameliorated site conditions compared to adjacent grasslands, although natural stands did so more effectively than managed ones. At the same time, all forests desiccated deeper soil layers during the vegetation period, and the longer the canopy lifetime, the more severe the desiccation in summer and more delayed the recharge after the active period of the canopy. We conclude that the site-scale environmental amelioration brought about by woody species is bound to co-occur with the desiccation of deeper soil layers, leading to deficient ground water recharge. This means that the cost of creating islands of fertility for sensitive herb layer organisms is an inevitable negative impact on regional water balance. The canopy type or management intensity of the forests affects the magnitude but not the direction of these effects. The outlined framework of the effects of woody species should be considered for the conservation, restoration, or profit-oriented use of forests as well as in forest-based carbon sequestration and soil erosion control projects in water-limited ecosystems.

## Introduction

Forest-grassland mosaics occur all over the world at the interfaces of forested and grassy biomes and take up various forms, such as forest-steppes and savannas (Scholes and Archer 1997, Erdős et al. 2018a), wood-pastures created by forest thinning and subsequent grazing (Bergmeier et al. 2010, Hartel et al. 2015), grasslands recently encroached by woody species (Van Auken 2009, Grellier et al. 2013), and partially afforested drylands, where trees are planted to reduce soil erosion, or to increase atmospheric carbon sequestration (Yosef et al. 2018, Bastin et al. 2019). Being prime ecosystem engineers, woody species profoundly affect biodiversity and ecosystem functions, including services for human populations. Due to the high global abundance and extreme diversity of forest-grassland ecosystems, there is an ongoing debate concerning the net effects of woody species, namely whether woody species promote or mitigate overall land degradation caused by local anthropogenic effects or even global environmental changes (Eldridge et al. 2011, Bastin et al. 2019, Lewis et al. 2019a, 2019b).

Woody species in arid and semi-arid ecosystems affect their surroundings on multiple scales (Scholes and Archer 1997). Locally, they moderate temperature extremes, particularly under the canopy, as shading leads to lower diurnal soil surface temperatures (Vetaas 1992). The canopy also reduces night-time radiative cooling compared to adjacent grassland sites (D’Odorico et al. 2013). Furthermore, woody species increase nutrient input under their canopy through litter turnover and by trapping and depositing airborne nutrient particles (Belsky 1994, Gea-Izquierdo et al. 2009). The resulting improved soil conditions, such as higher macroporosity, lead to higher water holding capacity (Cubera and Moreno 2007), which, coupled with lower diurnal temperatures, can reduce evaporative moisture loss (Maestre et al. 2003). In addition, deep-rooted woody species can passively translocate water from deeper moist zones to the topsoil by hydraulic lift (Yu and D’Odorico 2015), further ameliorating below-canopy conditions. However, the local effects of woody species on the moisture balance of the topsoil also have negative components. For instance, the canopy can intercept significant proportions of precipitation, reducing the amount of water reaching the soil (Farley et al. 2005). As a result, net effects of woody species on the moisture balance of the understory can range from positive (e.g. D’Odorico et al. 2007) to negative (e.g. Ludwig et al. 2004); yet more studies have reported positive effects in arid and semiarid ecosystems, contributing to the common view that solitary woody plants or clumps of them represent islands of fertility for understory organisms (Charley and West 1975, Schlesinger et al. 1990, Scholes and Archer 1997, Cable et al. 2012).

However, there is evidence that the regional water balance and productivity tend to be negatively affected by increasing tree cover in water-limited areas worldwide (Jackson et al. 2000, Farley et al. 2005, Zeng et al. 2009). This has been shown by decreases in the measures of regional water yield such as stream flows (Scott and Lesch 1997, Buytert et al. 2007) and groundwater levels (Jobbágy et al. 2004, Adane et al. 2018).

These apparently opposing views on the environmental effects of woody species have rarely been linked together into a multi-scale framework. Local effects of trees are mostly studied in natural or semi-natural ecosystems, such as savannas (e.g. Vetaas 1992, Ludwig et al. 2004), Mediterranean open woodlands (e.g. Cubera and Moreno 2007, Gea-Izquierdo et al. 2009) or continental forest-steppes (e.g. Tölgyesi et al. 2018), where native species constitute forest-grassland mosaic habitats. Regional scale studies, in contrast, frequently focus on areas actively afforested by non-native trees, with canopy and root traits different from those of native trees (e.g. Bosch and Hewlett 1982, Farley et al. 2005, Nosetto et al. 2005, Buytert et al. 2007, Cao et al. 2007). Consequently, the aims and scopes of local and regional scale studies rarely meet, hindering progress towards a multi-scale understanding of the effects of trees. It is thus unclear whether the different effects of woody species at different scales are bound to occur in the same systems at the same time, or do not necessarily co-occur.

In this study, we aimed to link the local and regional scale effects of woody species. We studied the Kiskunság Sand Ridge in central Hungary as a model system, because it is a semiarid region (Molnár et al. 2003, Kelemen et al. 2015), where the potential natural vegetation is forest-steppe (Erdős et al. 2018a, 2018b). Recent large scale afforestation has increased the cover of trees with canopy types significantly differing from native trees (Biró et al. 2008), providing the opportunity to test how canopy type modifies the local and regional effects of woody vegetation.

We hypothesised that the link between the local and regional scale effects of woody species can be best established by capturing processes in lower soil layers. Despite being rarely considered in studies dealing with the effects of woody species (but see e.g., Wang et al. 2018), lower soil layers can be assessed in a spatially explicit, fine scale manner, i.e. processes can be directly linked to the vegetation patches above. Lower soil layers are also directly associated with ground water recharge (Rimon et al. 2007, Adane et al. 2018) and thus with regional water balance and productivity (Fig. 1). Hence, we use the difference in lower soil moisture compared to adjacent grasslands as a measure of the effects of woody species on regional water balance.

**Figure 1.**
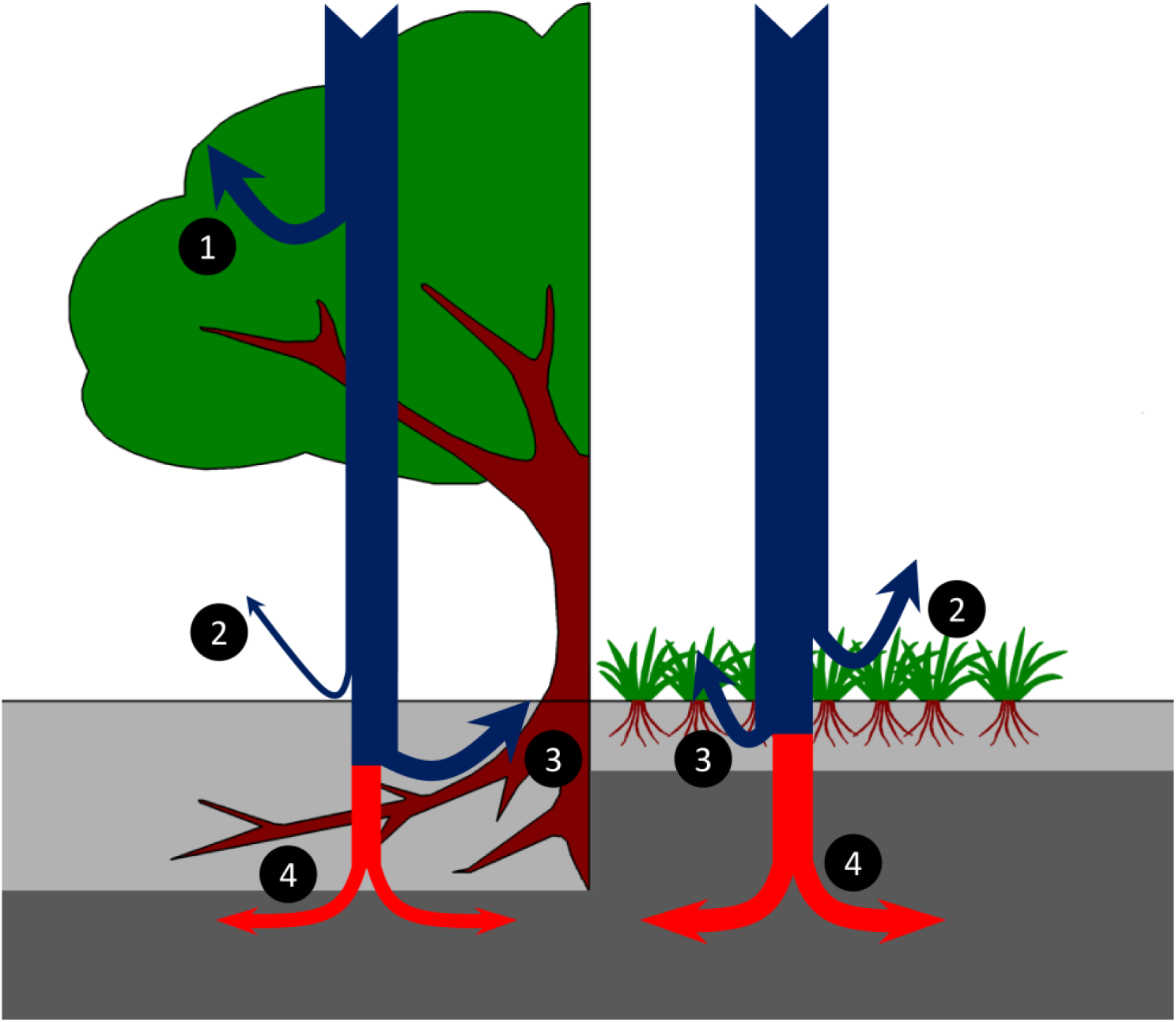
Interaction of woody and grassy vegetation with precipitation (blue arrows). The canopy of woody species intercepts an appreciable amount of precipitation (1), while, in grasslands, precipitation reaches the herb layer unhindered. Some moisture evaporates from the soil surface and the herb layer (2). Certain amounts of moisture seeping into the soil are absorbed by the roots (3). The proportion percolating through the rhizosphere (red arrows) decouples from the local vegetation and contributes to the regional water budget by eventually entering the groundwater. We assume the moisture contents of the lower parts of the rhizosphere and the layers below are indicative of the regional water balance by determining its input rate. Some woody species, can also tap into groundwater (not shown in the figure), also affecting the output rate, while grassy vegetation rarely does. Light grey layer: zone of the local effects of the vegetation; dark grey layer: zone of regional importance, i.e. the sub-rhizosphere vadose zone and the phreatic zone.

Specifically, we tested the following hypotheses: (1) Understory sites provide benign environmental conditions, i.e. reduced microclimatic extremes, moister topsoil and higher soil nutrient content compared to dry grassland sites (i.e. they function as “fertility islands”); (2) local effects of woody species are species specific and are mostly driven by differences in canopy types (annual canopy lifetime in particular); (3) woody vegetation reduces the fraction of precipitation percolating into deeper soil layers more than grassy vegetation but, outside vegetation period, soil moisture reserves recharge, and (4) canopy type substantially determines the dynamics of the desiccation–recharge cycles of deeper soil layers.

## Material and methods

### Study area and sampling sites

The study was performed in the Kiskunság Sand Ridge in central Hungary. The climate is continental with sub-Mediterranean influence. Annual precipitation is 550-600 mm and mean annual temperature is 11-12°C. The soil is coarse grained calcareous arenosol with low organic matter content. Groundwater is usually more than 3 meters below the soil surface (Tölgyesi et al. 2015). The potential vegetation is forest-steppe, i.e. a stable or slowly shifting mosaic of forest patches and grasslands (Erdős et al. 2015, 2018a). Natural forest patches are made up of *Populus alba* clones (hereafter poplar), while grasslands are bunch-grass steppes dominated by *Stipa* and *Festuca* species. Large areas of the natural vegetation have been cleared and turned into either low-productivity croplands or, more commonly, into plantations of drought tolerant, non-native trees (Biró et al. 2013). The most common planted trees include *Robinia pseudoacacia* (hereafter *Robinia*) and *Pinus nigra* (hereafter pine).

We selected four localities, where minimum 30-year-old, homogeneous stands of the three main types of trees, i.e. pine (evergreen), poplar (deciduous with long canopy lifetime) and *Robinia* (deciduous with short canopy lifetime) could be found close to each other at a similar elevation with grasslands available nearby (Fig. 2). The localities were as follows: Méntelek (N46.987, E19.571, 132 m a.s.l.), Fülöpháza (N46.861, E19.470, 109 m a.s.l.), Ágasegyháza (N46.804, E19.470, 104 m a.s.l.) and Izsák (N46.781, E19.326, 98 m a.s.l.). The distance between localities was 16.6±8.0 km (mean±SD). The average soil grain size was around 0.25 mm both in the top and deep layers in all habitats of every location; the clay fraction (<0.02 mm) was very low (3.67±1.42 m/m%; N=96).

**Figure 2.**
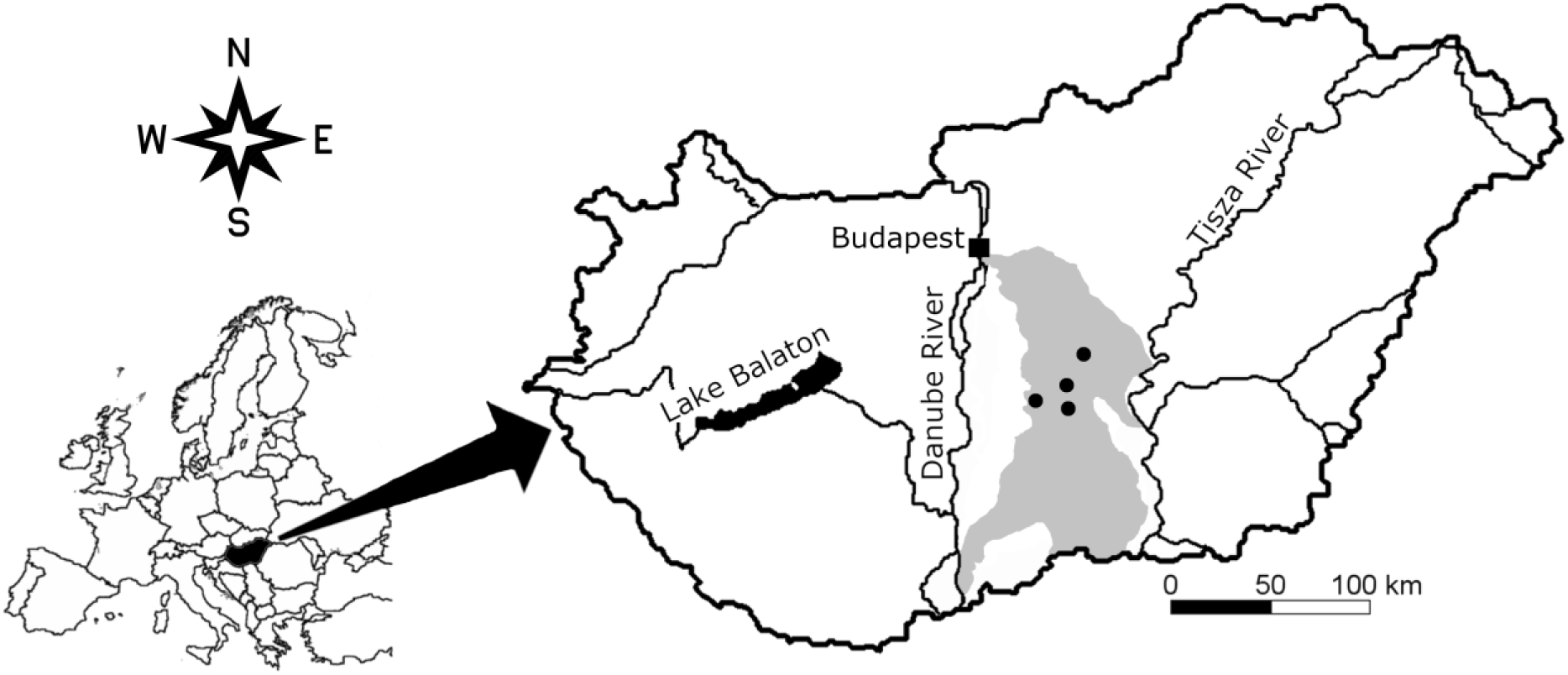
Geographical position of the studied localities (black dots) in the Kiskunság Sand Ridge (grey area), central Hungary.

### Sampling design

In each locality, we selected one stand for each forest type (poplar, *Robinia* and pine) and two patches of grasslands for sampling in 2017 and 2018. Forest stands ranged from 0.1 to 3 ha in size, while grasslands formed a more continuous matrix among forested patches. Grasslands were overrepresented to gain a more solid baseline to which we could compare the parameters of forested habitats. In every habitat patch of each locality (5 habitat patches × 4 localities), we measured microclimate (temperature and relative humidity) for 24-hour periods once every 6-7 weeks over the course of an entire year (measurement occasions: March 11-13, April 21-23, May 29-30, July 15-16, August 28-30, October 13-14, December 1-3 and January 20-21). We used Voltcraft DL-121TH data loggers installed 2-3 cm above ground and covered them by plastic roofs to avoid direct sunlight and to reduce frost precipitation on the sensors. In January, some sensors were fully covered in snow, while others could still communicate with above-snow air. This introduced a high variability in the data, confounding the effects of vegetation cover; therefore, we discarded this occasion from the analysis of microclimatic data.

To assess canopy cover, we took two to four digital photos of the canopy of each forest patch from 1 m above ground on every sampling occasion. According to Paletto and Tosi (2009), wide-angle photos overestimate canopy cover; therefore, we used a comparatively narrow angle setting (72 mm, equalling a diagonal angle of 34.4°). We took the photos near the microclimate stations (0-10 m from them) in spots where the trees had typical canopy structure and stem density. In October, we also took soil samples from the upper and lower soil layers of all habitat patches for soil analysis. The top 10 cm of the soil was too heterogeneous owing to a high amount of partially degraded litter and the thick rhizosphere of herbaceous species; therefore, we collected the upper sample from 10-20 cm deep. Since we encountered very few tree roots below 60 cm, we took the lower sample from 70-80 cm deep, using two replicates per patch (5 habitat patches × 4 localities × 2 replicates per patch, making 40 sampling sites) and measured humus and nitrogen content in a soil laboratory. Replicates within patches were spaced at least 20 m apart, in order to capture different tree individuals.

For soil moisture measurements, we drilled a 110 cm deep, cylindrical hole, with a diameter of 15 cm, in all the 40 sampling sites on each of the eight sampling occasions, using a soil auger. Roots that could not be tackled by the auger were gently cut through by a branch cutter. If a major root was encountered, we discarded the hole and started drilling another one nearby. We measured moisture content in every 10 cm soil depth segment as we proceeded downwards. We used a TDR 300 soil moisture meter, and made three vertical measurements in every segment, leading to a grand total of 11,520 soil moisture records covering the upper 120 cm of the soil. We considered 120 cm sufficiently deep because poplar and *Robinia* have the highest root density at soil depths of 20-80 cm and 20-60 cm, respectively (Cao et al. 2007) and pine has the majority of its roots, including fine roots, in the upper 50 cm (Hoffmann and Usoltsev 2001).

### Data analysis

We converted the canopy photos to black-and-white with manual thresholding (GIMP 2.8.10 software). Black pixels corresponded to leaves and branches and white pixels to the sky. We expressed canopy cover as the percentage of black pixels and used it as a proxy for the capacity of the canopy layer to intercept precipitation and its transpiration potential. Resulting canopy cover scores were compared across forest types with linear mixed-effects models. We built a model for each occasion and used locality as the random factor.

Regarding microclimatic recordings, synchronised 24-h periods could not be acquired for all five habitats of all four localities due to damage to the measuring stations (vandalism or damage by wild animals), and one of the localities provided reliable data only in two occasions (Table S1). Nevertheless, the resulting sample size per forest type (N=19-21) was suitable for statistical analysis. We standardized microclimatic data to the local average grassland values, i.e. average grassland data of a location were subtracted from the data of the adjacent forested habitats for every measurement time point. In the subsequent analysis, we separated daytime (10 a.m. – 4 p.m.) and night-time (10 p.m. – 4 a.m.) periods and avoided transitional periods during sunrise and sunset. We tested the effect of canopy cover on average daytime and night-time microclimatic variables (daytime and night-time temperature and relative humidity differences) with simple linear regression in the case of *Robinia* and poplar forests. The canopy of pine forests remained stable throughout the year and thus no cover gradient was available. Therefore we used one-sample t-tests with 0 as the hypothetical value to test whether the canopy of pine forests affects the microclimate compared to grasslands.

We prepared linear mixed-effects models for the soil parameters (humus and nitrogen content). We used habitat type (four levels: poplar, *Robinia*, pine and grassland) and depth layer (two levels: upper and lower) as ‘fixed-factors’ and included locality as ‘random factor’.

We averaged the three moisture values per depth segment of each hole and then grouped the 12 segments into six thicker layers (0-20 cm, 20-40 cm, etc.) by averaging the corresponding average records. Resulting moisture values were analysed with linear mixed-effects models (fixed effect: habitat type; random effect: locality). Separate models were prepared for each depth in each measurement occasion, resulting in a total of 48 models (8 occasions × 6 depth layers).

Statistical analysis was performed in R environment (R Core Team 2018). We used the ‘lme’ function of the *nlme* package for preparing linear mixed-effects models (Pinheiro et al. 2018). We used the built-in ‘anova’ function to test whether the models explain a significant proportion of the variation of the data. Pairwise comparisons of the levels of fixed factors with more than two levels (forest types and habitat types) were also considered in significant models. We adjusted the resulting pair-wise p-values for multiple comparisons using the fdr (false discovery rate) method (Benjamini and Hochberg 1995).

## Results

### Canopy cover

Canopy cover in pine forests was stable throughout the year (79.9±3.2%, mean±SD). By April, poplar forests had unfolded their leaves and their cover reached that of the pine forests. *Robinia* forests were still mostly dormant in April. In May, we detected similar levels of canopy cover in every forest type. Scores were similar in July, August and October. In December, as deciduous trees shed their leaves, their canopy cover dropped and remained low also in January (Fig. 3, Table S2).

**Figure 3.**
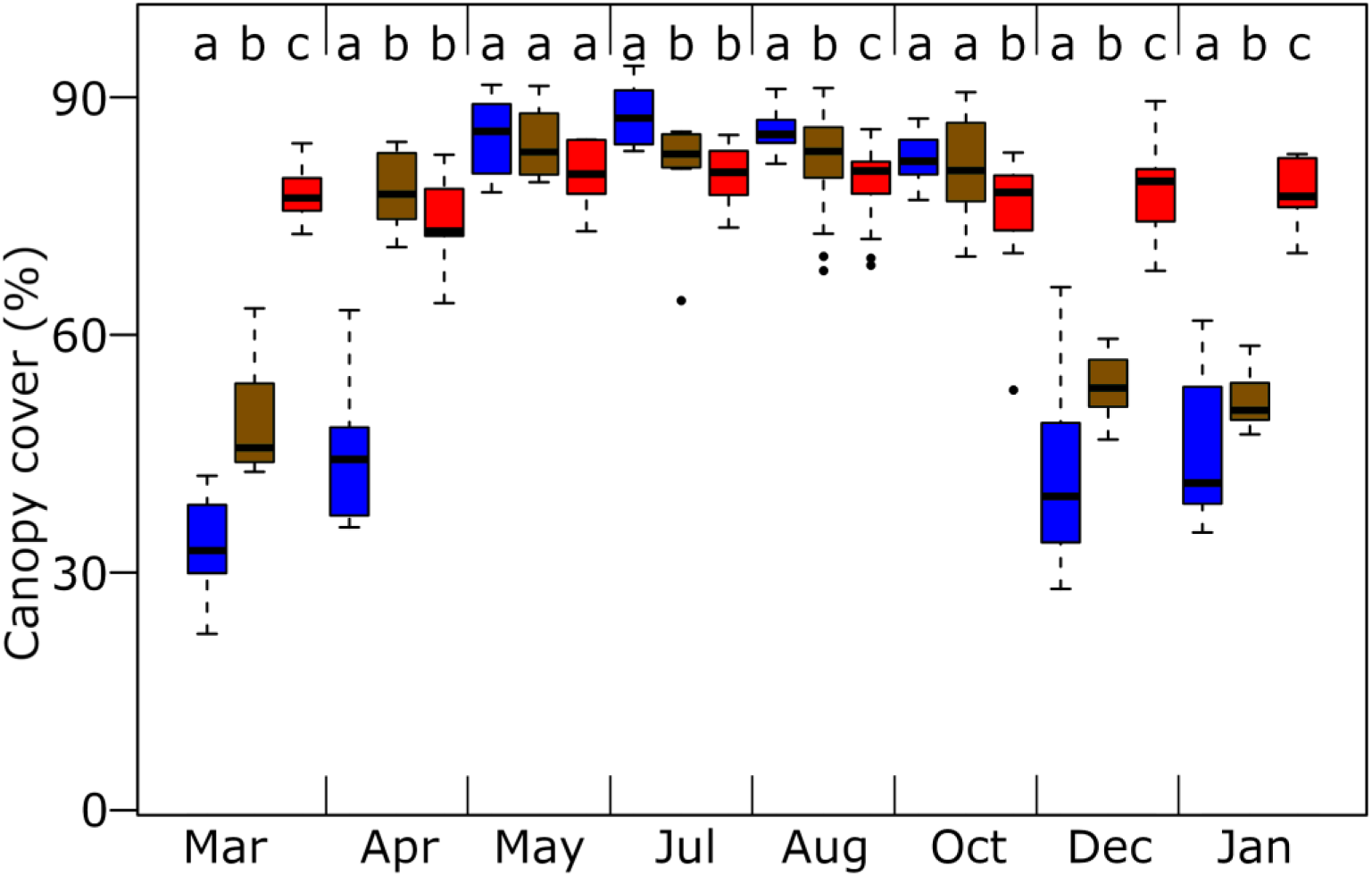
Canopy cover of *Robinia* (blue), poplar (green) and pine (red) forests over the course of a year. Lower case letters above boxes identify significantly different groups within each occasion (linear mixed-effects models).

**Figure 4.**
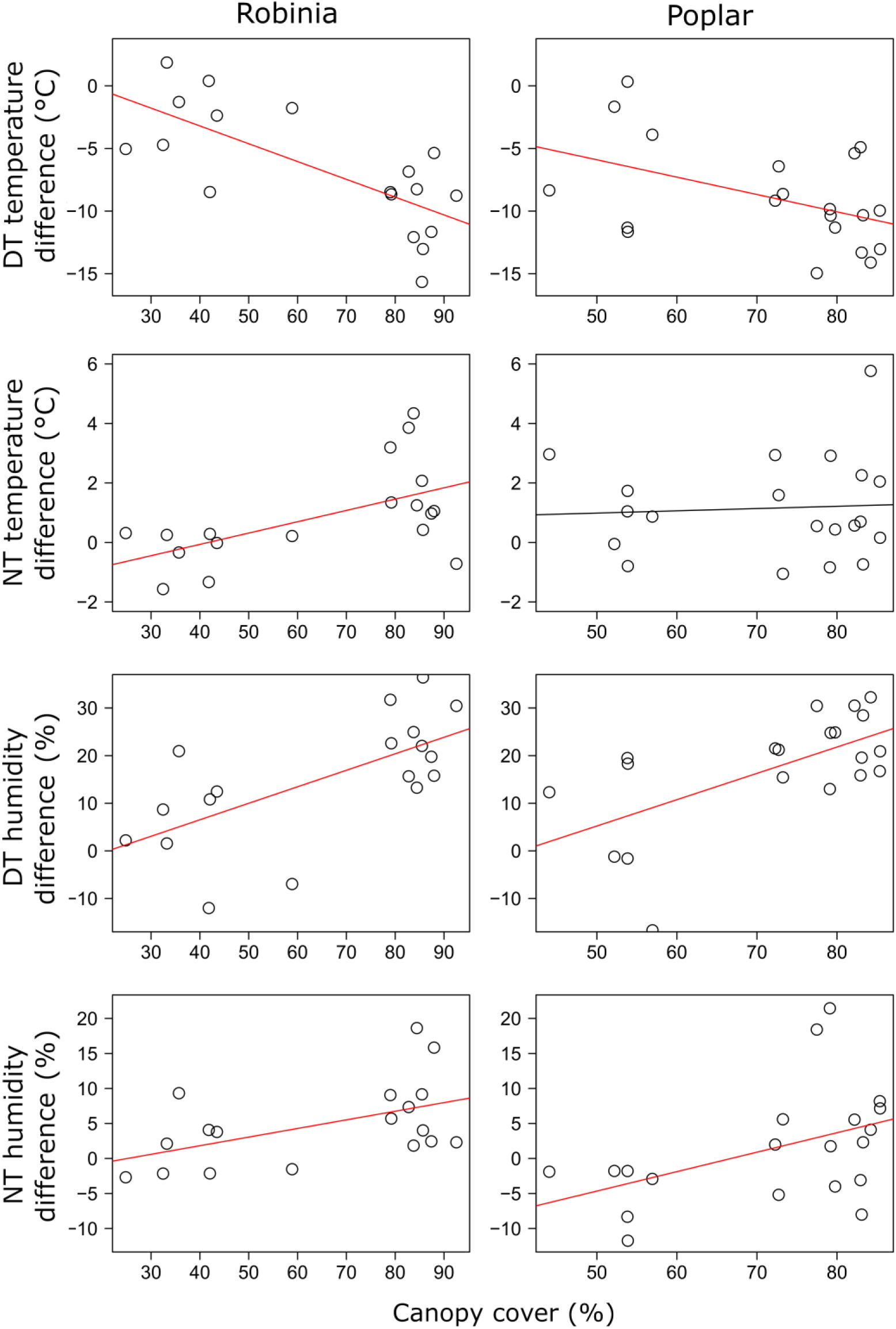
Effects of canopy cover on microclimatic variables in *Robinia* and poplar forests as compared to open grasslands. Pine forests are not shown because their canopy cover remained within a very narrow range throughout the year. Trend lines in red indicate significant relationship. DT: daytime; NT: night-time.

### Microclimate

Increasing canopy cover in *Robinia* forests decreased daytime but increased night-time temperature relative to grasslands. Humidity was affected positively by the canopy cover both during the day and at night, but the effect was stronger during the day, indicated by a more than two times higher estimate for daytime relationship than for the night-time one. Poplar forests showed the same pattern with the exception that no statistically significant increasing trend could be confirmed for night-time temperature (Fig. 3, Table S3 and Fig. S1-2). Under the stable canopy of pine forests, temperature was around 8.2±3.9°C lower during the day than outside, while night-time temperature was only slightly higher under the canopy. No statistically significant effect could be found for night-time humidity but daytime humidity was 14.0±9.7% higher than in the grasslands (Table S3, Fig. S1-2).

### Soil composition

Humus and nitrogen contents were higher in the upper layer of the soil than in the lower layer in all habitat types. Poplar forests had higher humus content than the other habitat types in both layers; *Robinia* forests, pine forests and grasslands had identically low humus contents. Nitrogen content was very high in *Robinia* forests, while the other habitat types had identically low values (Fig. 5, Table S4).

**Figure 5.**
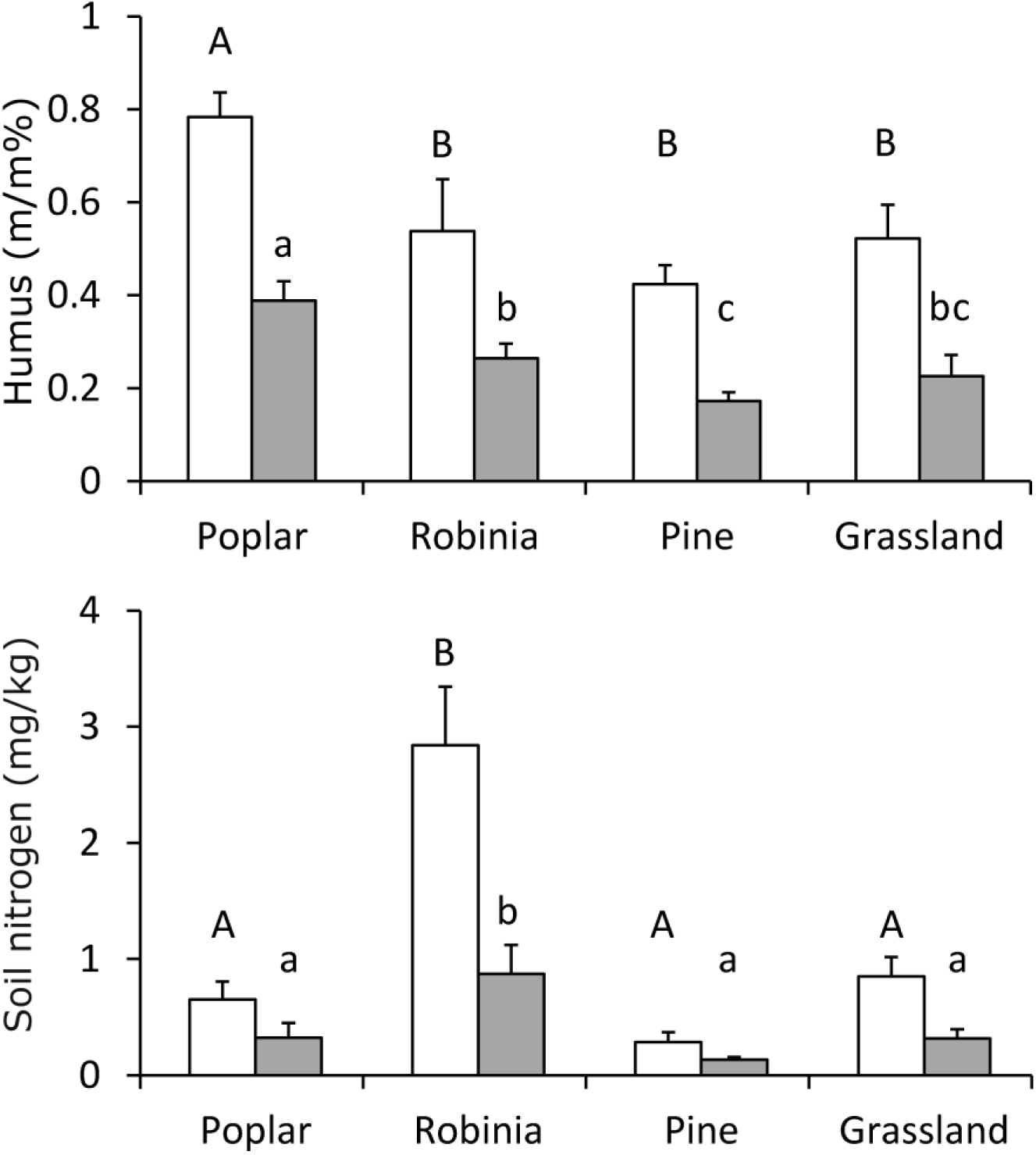
Humus and nitrogen content of the soil of sandy poplar, *Robinia* and pine forests and grasslands in the upper (10-20 cm, white bars) and lower soil layers (70-80 cm, grey bars). Different letters indicate significantly different groups. Capital and lower case letters belong to the upper and lower soil layers, respectively.

### Soil moisture

We found striking differences in the annual vertical soil moisture cycles of the studied habitat types (Fig. 6, Table S5). In March, poplar and *Robinia* forests had moister topsoil (upper 20 cm) than grasslands, but pine forests did not. At deeper layers, grasslands, poplar forests and *Robinia* forests had similar moisture contents, while pine forests were drier than the other habitats at depths of 40 and 60 cm. In April, all forest types were moister in the topsoil than grasslands. Poplar forests developed a dry zone at 100–120 cm deep, and, below 20 cm, pine forests depleted most of their soil moisture and became drier than any of the other habitats. In May, topsoil was moister in poplar and *Robinia* forests than in grasslands but the moisture content of the deeper soil layers of the forests remained below that of the grasslands. Pine forests were drier than the other habitats throughout the entire vertical soil section. In July, conditions were similar to May, although desiccation in the deeper soil layers of poplar forests progressed and the moisture contents became similar to those in pine forests. Topsoil conditions remained unchanged in August compared to May and July but by this time even *Robinia* forests had reached the same low moisture contents in most of the deeper soil layers as poplar and pine forests. The topsoil of poplar and *Robinia* forests was moister than that of grasslands in October but deeper soil layers started to recharge. Recharging of deeper layers showed a more advanced stage in December, with only pine forests being drier than grasslands. Recharge of deeper soil layers had been complete by January, with very similar moisture levels in the studied habitats along the soil section. The moisture content of deeper soil layers was between 5 and 6 v/v%, which was presumably the field capacity of the studied sand soil.

**Figure 6.**
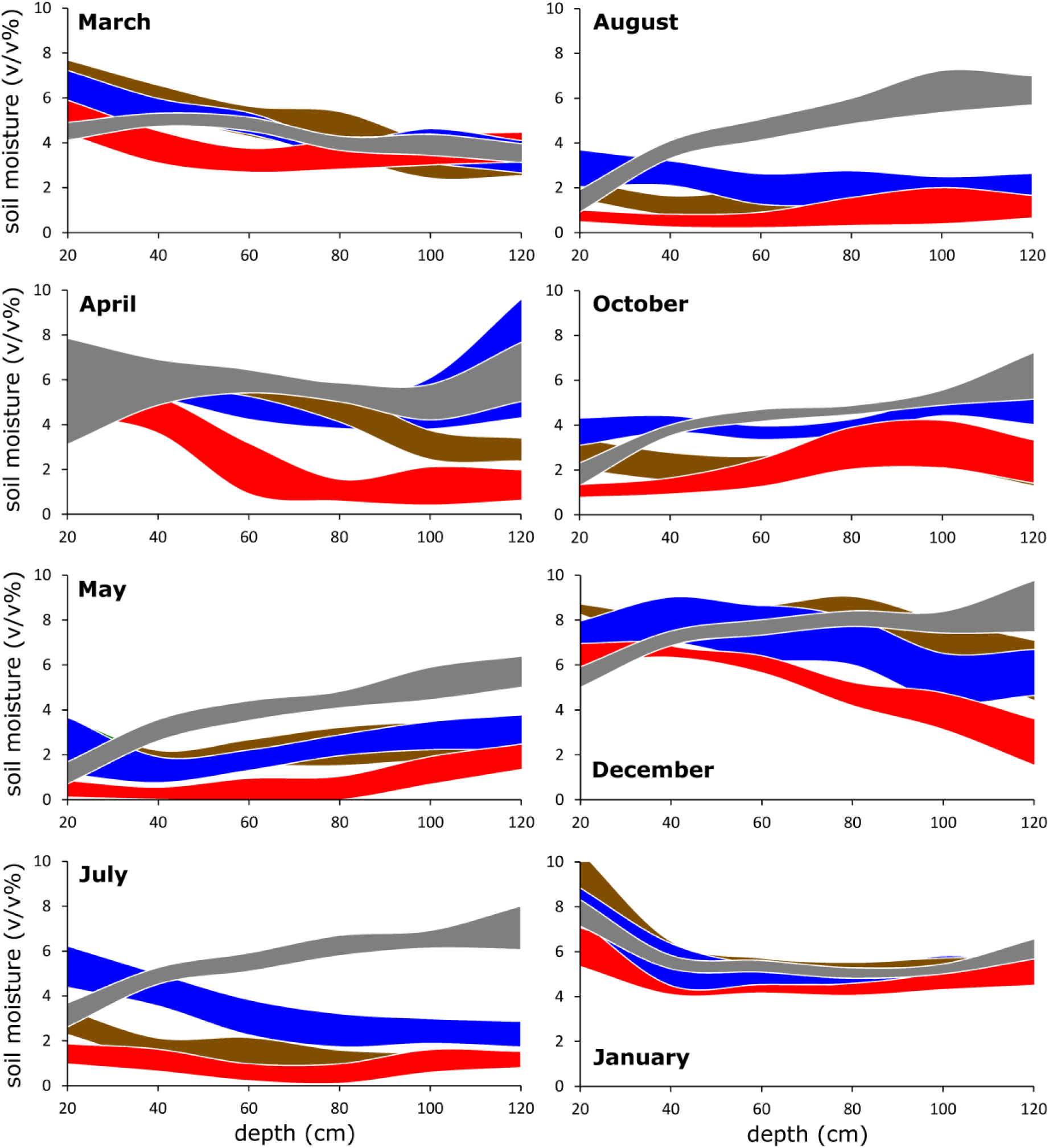
Vertical soil moisture distributions of sandy grasslands (grey), and poplar (green), *Robinia* (blue) and pine (red) forests in the upper 120 cm of the soil over the course of a year. Stripes encompass smoothed 95% confidence intervals. Stripes are over-plotted hierarchically (grassland > pine > *Robinia* > poplar).

## Discussion

### Islands of fertility?

Being prime ecosystem engineers, woody species profoundly modify local environmental parameters. Consequently, in mosaic habitats, where forested and grassy communities intermingle, trees introduce significant heterogeneity into the ecosystem (Fernandez-Moya et al. 2011, López-Sanchez et al. 2016). In the studied semiarid ecosystem, we found evidence for this environmental heterogeneity in microclimate, soil nutrient content and topsoil moisture. Organisms inhabiting the understory of the studied forest patches do not need to face the diurnal heat stress typical of grasslands in summer, and the cold stress is also reduced during spring nights, which is a sensitive period for seedlings and young shoots of understory plants (Coop et al. 2008, Inouya 2008). In addition, from spring till autumn, both air humidity and the topsoil moisture tended to be higher under the canopy of trees than in the grasslands, further ameliorating conditions for drought-intolerant organisms. We also found that poplar forests accumulated more humus in both studied soil layers than grasslands, while nitrogen was more abundant in *Robinia* forests than in grasslands, due to the high nitrogen-fixing capacity of this species (Cierjacks et al. 2013, Vítková et al. 2017). Our data thus confirm that the forest patches of the studied region represent islands of fertility; therefore, the studied forest-grassland mosaic appears typical of arid and semiarid ecosystems, with respect to the local effects of woody species (Charley and West 1975, Schlesinger et al. 1990, Scholes and Archer 1997, Cable et al. 2012).

### Canopy type and micro-site effects

The effects of the different tree species on understory conditions can be associated to their canopy types, especially annual canopy lifetime. Since the canopy cover of *Robinia* forests is low during spring, they cannot protect the understory from occasional night frost as well as pine or poplar trees. In addition, poplar and pine forests can provide shelter for early spring, cool-adapted organisms during the day. In temperate forest–grassland mosaics such as the forest-steppe zone, these effects may be important ecosystem functions of tree canopies; however, *Robinia* forests cannot perform them.

We found that poplar forests increased the humus content in the topsoil compared to grasslands, but *Robinia* and pine forests did not. This can be associated to management intensity, as poplar forests are unmanaged natural habitats, while *Robinia* and pine stands are plantations exposed to periodic soil disturbance during forestry activities (Zeng et al. 2009). This effect outweighed the high nitrogen content in *Robinia* forests, which could otherwise promote high herb layer productivity, but here was not enough to support higher humus formation in the soil. The lack of increased humus content in *Robinia* forests can also be explained by the low sclerenchyma and overall organic matter content of the leaves per area unit, which may also enable quicker decomposition. These assumptions are supported by the high specific leaf area of *Robinia* (*Robinia pseudoacacia*, 17.9 mm^2^ mg^-1^) compared with poplar (*Populus alba*, 11.2 mm^2^ mg^-1^) and pine (*Pinus sylvestris*, 5.4 mm^2^mg^-1^; source: LEDA trait base; Kleyer et al. 2008).

Additionally, the topsoil of pine forests was drier than that of grasslands in summer. This can be caused by a more superficial root system but the most likely explanation is the high canopy interception of precipitation in pine forests. Although the canopy cover showed little variation between forest types in summer, there is evidence that the thick bunches of needles in the canopy of coniferous trees intercept more precipitation than the flat leaves of deciduous trees (Silva and Rodríguez 2001; Farley et al. 2005; Brygadyrenko 2014). This effect seems to hinder the creation of moist enclaves in the understory of pine trees in semiarid ecosystems during summer (see also Tölgyesi et al. 2018, for similar findings in central Asia). Natural stands of poplar trees therefore represented the highest quality islands of fertility, compared to the managed plantations of the other tree species.

### Lower soil moisture

In summer, when water balance was lowest due to high diurnal temperatures and high rates of transpiration, all forest types removed most of the moisture content from lower soil layers. At the same time, grasslands did not tap the moisture stored in lower soil layers; moisture content in grasslands remained constant below 40 cm throughout the year. The approximately 5 v/v% we detected is likely to be the field capacity of the coarse-grained sand soils of the studied ecosystem.

The fact that soil moisture was periodically below field capacity in the deeper forest soil layers suggests forest soils experience less groundwater recharge following precipitation than grasslands, a phenomenon which has been observed elsewhere (e.g. Walter and Breckle 1986). During these periods, the topsoil of grasslands was also very dry, but the thickness of this top layer was a fraction of the dry layer we found under the trees. Our findings thus demonstrate how individual trees as well as forests and tree plantations may contribute to decreases of water balance at the regional scale. Groundwater in the local Kiskunság Sand Ridge dropped by 1-3 m on average in the second half of the 20^th^ century (Szilágyi and Vörösmarty 1997) and intensive afforestation was suspected as a major driver (Zsákovics et al. 2007), although the extreme desiccating effect of trees in lower soil layers was hitherto unknown. Our study provides spatially explicit evidence for this effect which is coherent within the context of regional moisture dynamics.

Interestingly, the observed below-ground moisture pattern is little recorded in the international scientific literature, for which we posit two possible explanations. First, this pattern may be specific to the studied region; second, it is more common but mostly overlooked. We deem the first explanation unlikely because our study confirmed the island of fertility concept, which is typical for semiarid forest-grassland mosaics; therefore, there is little indication that untypical processes should take place below the topsoil. However, in finer textured soils, which have different water holding capacity and thus different rates of infiltration, the pattern may be less pronounced or even different. For instance, Warren et al. (2005) showed that lower soil layers can be moister than the topsoil both in semiarid grasslands and pine forests, although those lower soil layers had finer texture than the topsoil. The observed pattern may also differ in extremely arid areas which lack enough precipitation to infiltrate lower layers. For instance, Bhark and Small (2003) showed that water infiltration rarely exceeds half a meter after rainy periods in woody–grassy mosaics of New Mexico, while Yang et al. (2012) also found a lack of deep soil recharge in arid forest-steppes of the Chinese Loess Plateau. Nevertheless, large sandy regions with sparse tree cover and good soil infiltration capacity, such as the Namibrand in South Africa (Krug 2017), the Keerqin area in NE China (Zeng et al. 2009), the Nebraska Sand Hills in North America (Adane et al. 2018) or the Doñana sand dunes in Spain (Munoz-Reinoso and Novo 2005) may show soil moisture patterns similar to our findings.

The second possible explanation for the lack of reported different effects of trees on topsoil and lower soil layers may be the frequently low resolution of layer thickness. For instance, Yang et al. (2012) studied the upper 8 m of the soil under afforested areas in China but only considered 1 m thick layers. Although Cubera and Moreno (2007) found that deep soil layers (1–2 m) were moister below grasslands than below trees in a Mediterranean open woodland, they found no clear trend for the upper 1 m and did not investigate differences between thinner layers of soil (<1m). Thus, we believe the soil moisture pattern we described may be widespread, but can only be detected with a high-resolution analysis of the soil layers. Nevertheless, further research is needed, under different macroclimatic conditions and soil textures, to identify drivers of desiccation of lower soil layers while maintaining improved microsite conditions in the understory.

### Canopy type and lower soil moisture

Canopy type had a clear effect on the desiccation kinetics of lower soil layers. The earliest seasonal decline in soil moisture was detected under pine forests, corresponding to their high canopy interception of precipitation throughout the year, as well as the ability to transpire during warm late winter and early spring days (Shulze et al. 2002). The canopy cover scores of poplar forests increased earlier than those of *Robinia* forests, which also correlated with the earlier decline of soil moisture under poplar forests than under *Robinia* plantations. At the end of the vegetation period, the recharge under deciduous forests could also occur faster than below pine forests, since deciduous trees lost most of their transpiration and interception capacity as they shed leaves, while interception did not change in pine forests. Our results thus suggest that the longer the annual lifetime of the canopy, the more intensive and more prolonged the desiccating effect on lower soil layers. This trend conforms to regional scale differences in water yield effects of different trees. For instance, *Populus tremula*, a close relative of the poplar species of the present study (*P. alba*), causes a 20-40% regional water deficit compared to grasslands in a forest-steppe area of the Chinese Loess Plateau (Cao et al. 2008), while water yield reductions after afforestation with pines amount for 40-50%, largely irrespective of macroclimatic conditions (Farley et al. 2005, Buytert et al. 2007). To the best of our knowledge, regional drying effects of deciduous forests with short and long canopy lifetime have not been previously compared, but based on our results, we can assume a lower regional water balance in areas with deciduous forests with longer annual canopy lifetime.

### Conclusions

Our findings revealed that trees in a semiarid, sandy forest–grassland ecosystem ameliorate micro-site conditions under their canopy compared to grasslands. This ‘island of fertility’ effect is stronger in natural forest stands, but intensively managed tree plantations also temper micro-environmental extremes for understory organisms. However, the cost of this local effect is the desiccation of lower soil layers in all forest types, leading to deficient groundwater recharge below forests. This finding establishes a direct link between the local ameliorating effects and the regional drying effects of trees, which, we conclude, can co-occur in the same ecosystems simultaneously. These concordant effects of woody species on different scales must be carefully considered during the conservation or restoration of natural forest–grassland mosaics, the profit-oriented use of tree plantations, and the afforestation of water-limited areas to target soil erosion control and, in particular, carbon sequestration. The potential of forest-based carbon sequestration to mitigate climate change has received great attention lately. The best-case scenario of Bastin et al. (2019), which calculates with the afforestation of all land that can maintain trees has promising results, while others raise serious concerns about the credibility of their model (e.g. Lewis et al. 2019a). For instance, afforestation entails a drop in albedo in high latitudes, which enhances warming (Lewis et al. 2019b). Veldman et al. (2019) argues against the afforestation of sparsely wooded ecosystems, such as savannas, as it would likely cause biodiversity loss. Our findings highlight that drylands with coarse soil texture are especially weak candidates for climate mitigation by afforestation, but if land managers decide for afforestation anyway, trees with short annual canopy lifetime should be chosen.

## Declarations (blinded)

### Data accessibility

Data used in this paper will be stored in the Dryad Digital Repository upon acceptance.

## Supplementary Information

**Table S1** Overview of the microclimatic measurements throughout the study.

**Table S2** Results of the linear mixed-effects models prepared for the canopy cover of sandy poplar, *Robinia* and pine forests.

**Table S3** Results of the linear models prepared for the effect of canopy cover on microclimatic variables in Robinia and poplar forests, and the one-sample t-tests on the effect of pine forests on microclimate as compared to adjacent open grasslands.

**Table S4** Results of the linear mixed-effects models prepared for humus and nitrogen contents of the soil of sandy grasslands and poplar, *Robinia* and pine forests.

**Table S5** Results of the linear mixed-effects models prepared for the soil moisture content of sandy grasslands and poplar, *Robinia* and pine forests over the course of a year.

## References

Adane, Z. A., Nasta, P., Zlotnik, V. and Wedin, D. 2018. Impact of grassland conversion to forest on groundwater recharge in the Nebraska Sand Hills. – J Hydrol: Reg Stud 15: 171–183.

Bastin, J.-F., Finegold, Y., Garcia, C., Mollicone, D., Rezende, M., Routh, D., Zohner, C. M. and Crowther, T. W, 2019. The global tree restoration potential. – Science 375: 76–79.

Belsky, A. J. 1994. Influences of trees on savanna productivity: Tests of shade, nutrients, and tree–grass competition. – Ecology 75: 922–932.

Benjamini, Y. and Hochberg, Y. 1995. Controlling the false discovery rate: a practical and powerful approach to multiple testing. – Journal of the Royal Statistical Society Series B 57: 289–300.

Bergmeier, E., Peterman, J. and Schröder, E. 2010. Geobotanical survey of wood-pasture habitats in Europe: diversity, threats and conservation. – Biodivers Conserv 19: 2995–3014.

Biró, M., Révész, A., Molnár, Z., Horváth, F. and Czúcz, B. 2008. Regional habitat pattern of the Danube-Tisza Interfluve in Hungary II. – Acta Bot Hung 50: 19–60.

Biró, M., Szitár, K., Horváth, F., Bagi, I. and Molnár, Z. 2013. Detection of long-term landscape changes and trajectories in a Pannonian sand region: comparing land-cover and habitat-based approaches at two spatial scales. – Community Ecol 14: 219–230.

Bhark, E. W. and Small, E. E. 2003. Association between plant canopy and the spatial patterns of infiltration in shrubland and grassland of the Chihuahuan Desert, New Mexico. – Ecosystems 6: 185–196.

Bosch, J. M. and Hewlett, J. D. 1982. A review of catchment experiments to determine the effects of vegetation changes on water yield and evaporation. – J Hydrol 55: 3–23.

Brygadyrenko, V. V. 2014. Influence of soil moisture on litter invertebrate community structure of pine forests of the steppe zone of Ukraine. – Folia Oecol 41: 8–16.

Buytaert, W., Iniguez, V. and De Bievre, B. 2007. The effects of afforestation and cultivation on water yield in the Andean páramo. – For Ecol Manag 251: 22–30.

Cable, J. M., Barron-Gafford, G. A., Ogle, K., Pavao-Zuckerman, M., Scott, R. L., Williams, D. G. and Huxman, T. E. 2012. Shrub encroachment alters sensitivity of soil respiration to temperature and moisture. – J Geophys Res 117: G01001.

Cao, S., Chen, L., Xu, C. and Liu, Z. 2007. Impact of three soil types on afforestation in China’s Loess Plateau: Growth and survival of six tree species and their effects on soil properties. – Landsc Urban Plan 83: 208–217.

Charley, J. L. and West, N. E. 1975. Plant-induced soil chemical patterns in some shrub-dominated semi-desert ecosystems of Utah. – J Ecol 63: 945–963.

Coop, J. D. and Givnish, T. J. 2008. Constraints on tree seedling establishment in montane grasslands of the Valles Caldera, New Mexico. – Ecology 89: 1101–1111.

Cierjacks, A., Kowarik, I., Joshi, J., Hempel, S., Ristow, M., von der Lippe, M. and Weber, E. 2013. Biological flora of the British Isles: Robinia pseudoacacia. – J Ecol 101: 1623–1640.

Cubera, E. and Moreno, G. 2007. Effect of land-use on soil water dynamic in dehesas of Central-Western Spain. – Catena 71: 298–308.

D’Odorico, P., Caylor, K., Okin, G. S. and Scanlon, T. M. 2007. On soil moisture–vegetation feedbacks and their possible effects on the dynamics of dryland ecosystems. – J Geophys Res 112: G04010.

D’Odorico, P., He, Y., Collins, S., De Wekker, S. F. J., Engel, V. and Fuentes, J. D. 2013. Vegetation-microclimate feedbacks in woodland-grassland ecotones. – Glob Ecol Biogeogr 22: 364–379.

Eldridge, D. J., Bowker, M. A., Maestre, F. T., Roger, E., Reynolds, J. F. and Whitford, W. G. 2011. Impacts of shrub encroachment on ecosystem structure and functioning: towards a global synthesis. – Ecol Lett 14: 709–722.

Erdős, L., Tölgyesi, C., Cseh, V., Tolnay, D., Cserhalmi, D., Körmöczi, L., Gellény, K. and Bátori, Z. 2015. Vegetation history, recent dynamics and future prospects of a Hungarian sandy forest-steppe reserve: forest-grassland relations, tree species composition and size-class distribution. – Community Ecol 16: 95–105.

Erdős, L., Ambarli, D., Anenkhonov, O. A., Bátori, Z., Cserhalmi, D., Kiss, M., Kröel-Dulay, G., Liu, H., Magnes, M., Molnár, Z., Naqinezhad, A., Semenishchenkov, Y. A., Tölgyesi, C. and Török, P. 2018a. The edge of two worlds: A new synthesis on Eurasian forest-steppes. – Appl Veg Sci 21: 345–362.

Erdős, L., Kröel-Dulay, G., Bátori, Z., Kovács, B., Németh, C., Kiss, P. J. and Tölgyesi, C. 2018b. Habitat heterogeneity as a key to high conservation value in forest-grassland mosaics. – Biol Conserv 226: 72–80.

Farley, K. A., Jobbágy, E. G. and Jackson, R. B. 2005 Effects of afforestation on water yield: a global synthesis with implication for policy. – Glob Change Biol 11: 1565–1576.

Fernández-Moya, J., San Miguel-Ayanz, A., Canellas, I. and Gea-Izquierdo, G. 2011. Variability in Mediterranean annual grassland diversity driven by small-scale changes in fertility and radiation. – Plant Ecol 212: 865–877.

Gea-Izquierdo, G., Montero, G. and Canellas, I. 2009. Changes in limiting resources determine spatio-temporal variability in tree-grass interactions. – Agrofor Syst 76: 375–387.

Grellier, S., Ward, D., Janeau, J.-L., Podwojewski, P., Lorentz, S., Abbadie, L., Valentin, C. and Barot, S. 2013. Positive versus negative environmental impacts of tree encroachment in South Africa. – Acta Oecol 53: 1–10.

Hartel, T., Plieninger, T. and Varga, A. 2015. Wood-pastures of Europe. – In: Kirby, K. and Watkins, C. (eds.), Europe’s changing woods and forests: from wildwood to managed landscapes. CABI Press, pp. 61–76.

Hoffmann, C.W. and Usoltsev, V. A. 2001. Modelling root biomass distribution in Pinus sylvestris forests of the Turgai Depression of Kazakhstan. – For Ecol Manag 149: 103–114.

Inouya, D. W. 2008. Effects of climate change on phenology, frost damage, and floral abundance of montane wild flowers. – Ecology 89: 353–362.

Jackson, R. B., Schenk, H. J., Jobbágy, E. G., Canadell, J., Colello, G. D., Dickinson, R. E., Field, C. B., Friedlingstein, P., Heimann, M., Hibbard, K., Kicklighter, D. W., Kleidon, A., Neilson, R. P., Parton, W. J., Sala, O. E. and Sykes, M. T. 2000. Belowground consequences of vegetation change and their treatment in models. – Ecol Appl 10: 470–583.

Jobbágy, E. G. and Jackson, R. B. 2004. Groundwater use and salinization with grassland afforestation. – Glob Change Biol 10: 1299–1312.

Kelemen, A., Valkó, O., Kröel-Dulay, G., Deák, B., Török, P., Tóth, K., Miglécz, T. and Tóthmérész, B. 2016. The invasion of common milkweed (Asclepias syriaca L.) in sandy old-fields – Is it a threat to the native flora? – Appl Veg Sci 19: 218–224.

Kleyer, M., Bekker, R. M., Knevel, I. C., Bakker, J. P., Thompson, K., Sonnenschein, M., Poschlod, P., Van Groenendael, J. M., Klimeš, L., Klimešová, J., Klotz, S., Rusch, G. M., Hermy, M., Adriaens, D., Boedeltje, G., Bossuyt, B., Dannemann, A., Endels, P., Götzenberger, L., Hodgson, J. G., Jackel, A.-K., Kühn, I., Kunzmann, D., Ozinga, W. A., Römermann, C., Stadler, M., Schlegelmilch, J., Steendam, H. J., Tackenberg, O., Wilmann, B., Cornelissen, J. H. C., Eriksson, O., Garnier, E. and Peco B. 2008. The LEDA Traitbase: A database of life-history traits of northwest European flora. – J Ecol 96: 1266–1274.

Krug, J. H. A. 2017. Tree water potentials supporting an explanation for the occurrence of Vachellia erioloba in the Namib Desert (Namibia). – For Ecosyst 4: 20.

Lewis, S. L., Mitchard E. T. A., Prentice, C., Maslin, M. and Poulter, B. 2019a. Comment on “The global tree restoration potential”. – Science 366, eaaz0388, DOI: 10.1126/science.aaz0388

Lewis, S. L., Wheeler, C. E., Mitchard, E. T. A. and Koch, A. 2019b. Restoring natural forests is the best way to remove atmospheric carbon. – Nature 568: 25–29.

López-Sánchez, A., Miguel, A. S., López-Carrasco, C., Huntsinger, L. and Roig, S. 2016. The important role of scattered trees on the herbaceous diversity of grazed Mediterranean dehesa. – Acta Oecologica 76: 31–38.

Ludwig, F., de Kroon, H., Berendse, F. and Prins, H. H. T. 2004. The influence of savanna trees on nutrient, water and light availability and the understorey vegetation. – Plant Ecol 170: 93–105.

Maestre, F. T., Bautista, S. and Cortina, J. 2003. Positive, negative, and net effects in grass– shrub interactions in mediterranean semiarid grasslands. – Ecology 84: 3186–3197.

Molnár, Z., Sipos, F., Vidéki, R., Biró, M. and Iványosi-Szabó, A. 2003. Dry sand vegetation of the Kiskunság. TermészetBÚVÁR Alapítvány Kiadó.

Munoz-Reinoso, J. C. and Novo, F. G. 2005. Multiscale control of vegetation patterns: the case of Donana (SW Spain). – Landsc Ecol 20: 51–61.

Nosetto, M. D., Jobbágy, E. G. and Paruelo, J. M. 2005. Land-use change and water losses: the case of grassland afforestation across soil textural gradient in central Argentina. – Glob Change Biol 11: 1101–1117.

Paletto, A. and Tosi, V. 2009. Forest canopy cover and canopy closure: comparison of assessment techniques. – Eur J For Res 128: 265–272.

Pinheiro, J., Bates, D., DebRoy, S., Sarkar, D. and R Core Team 2019. nlme: Linear and Nonlinear Mixed Effects Models. R package version 3.1-142, https://CRAN.R-project.org/package=nlme.

R Core Team 2018. R: A language and environment for statistical computing. R Foundation for Statistical Computing, Vienna, URL https://www.R-project.org/

Rimon, Y., Dahan, O., Nativ, R. and Geyer, S. 2007. Water percolation through the deep vadose zone and groundwater recharge: Preliminary results based on a new vadose zone monitoring system. – Water Resource Res 43: W05402.

Schlesinger, W. H., Reynolds, J. F., Cunningham, G. L., Huenneke, L. F., Jarrell, W. M., Virginia, R. A. and Whithford, W. G. 1990. Biological feedbacks in global desertification. – Science 247: 1043–1048.

Scholes, R. J. and Archer, S. R. 1997. Tree-grass interactions in savannas. – Ann Rev Ecol Syst 28: 517–544.

Schulze, E.-D., Beck, E. and Müller-Hochenstein, K. 2002. Plant Ecology. – Springer-Verlag.

Scott, D. F. and Lesch, W. 1997. Streamflow responses to afforestation with Eucalyptus grandis and Pinus patula and to felling in the Mokobulaan experimental catchments, South Africa. – J Hydrol 199: 360–377.

Silva, I. C. and Rodríguez, H. G. 2001. Interception loss, throughfall and stemflow chemistry in pine and oak forests in northeastern Mexico. – Tree Physiol 21: 12–13.

Szilágyi, J. and Vörösmarty, C. 1997. Modelling unconfined aquifer level reductions in the area between the Danube and the Tisza Rivers in Hungary. – J Hydrol Hydromech 45: 328–347.

Tölgyesi, C., Zalatnai, M., Erdős, L., Bátori, Z., Hupp, N. R. and Körmöczi, L. 2015. Unexpected ecotone dynamics of a sand dune vegetation complex following water table decline. – J Plant Ecol 9: 40–50.

Tölgyesi, C., Valkó, O., Deák, B., Kelemen, A., Bragina, T. M., Gallé, R., Erdős, L. and Bátori, Z. 2018. Tree–herb co-existence and community assembly in forest-steppe transitions. – Plant Ecol Divers 11: 465–477.

Van Auken, O. W. 2009. Causes and consequences of woody plant encroachment into western North American grasslands. – J Environ Manag, 90: 2931–2942.

Veldman, J. W., Aleman, J. C., Alvarado, S. T., Anderson, T. M., Archibald, S., Bond, W. J., Boutton, T. W., Buchmann, N. et al. 2019. Comment on “The global tree restoration potential”. – Science 366, DOI: 10.1126/science.aay7976

Vetaas, O. R. 1992. Micro-site effects of trees and shrubs in dry savannas. – J Veg Sci 3: 337–344.

Vítková, M., Müllerová, J., Sádlo, J., Pergl, J. and Pysek, P. 2017. Black locust (Robinia pseudoacacia) beloved and despised: A story of an invasive tree in Central Europe. – For Ecol Manag 384: 287–302.

Wang, Z., He, Q., Hu, B., Pang, X. and Bao, W. 2018. Gap thinning improves soil water content, changes the vertical water distribution and decreases the fluctuation. – Can J For Res 48: 1042–1048.

Warren, J. M., Meinzer, F. C., Brooks, J. R. and Domec, J. C. 2005. Vertical stratification of soil water storage and release dynamics in Pacific Northwest coniferous forests. – Agric For Meteorol 130: 39–58.

Yang, L., Wei, W., Chen, L. and Mo, B. 2012. Response of deep soil moisture to land use and afforestation in the semi-arid Loess Plateau, China. – J Hydrol 475: 111–122.

Yosef, G., Walko, R., Avisar, R., Tatarinov, F., Rotenberg, E. and Yakir, D. 2018. Large-scale semi-arid afforestation can enhance precipitation and carbon sequestration potential. – Sci Rep 8:996.

Yu, K. and D’Odorico, P. 2015 Hydraulic lift as a determinant of tree–grass coexistence on savannas. – New Phytol 207: 1038–1051.

Zeng, D. H., Hu, Y. L., Chang, S. X. and Fan, Z. P. 2009. Land cover change effects on soil chemical and biological properties after planting Mongolian pine (Pinus sylvestris var. mongolica) in sandy lands in Keerqin, northeastern China. – Plant Soil 317: 121–133.

Zsákovics, G., Kovács, F., Kiss, A. and Pócsik, E. 2007. Risk analysis of the aridification-endangered sand-ridge area in the Danube-Tisza Interfluve. – Acta Climatol Chorol 40-41: 169–178.

