## Supplementary material for "Underground deserts below fertility islands? – Woody species desiccate lower soil layers in sandy drylands"

**Table S1** Overview of the microclimatic measurements throughout the study. Gr: grassland; Po: poplar forest; Ro: *Robinia* forest; Pi: pine forest; ✓: operational microclimate station; ×: malfuctioning or damaged microclimate station. There were two grassland stations in each locality, which is indicated by two functionality symbols in respective cells. If both grassland stations of a locality were compromised, we could not use otherwise operational forest stations of the same locality (also indicated with "×" sign), as we could not standardize them.

| | Fülöpháza | | | | Ágasegyháza | | | | Méntelek | | | | Izsák | | | |
|---|---|---|---|---|---|---|---|---|---|---|---|---|---|---|---|---|
| Month | Gr | Po | Ro | Pi | Gr | Po | Ro | Pi | Gr | Po | Ro | Pi | Gr | Po | Ro | Pi |
| March | ✓✓ | ✓ | ✓ | ✓ | ✓✓ | ✓ | ✓ | ✓ | ✓✓ | ✓ | ✓ | × | ×× | × | × | × |
| April | ✓✓ | ✓ | × | ✓ | ×× | × | × | × | ✓✓ | ✓ | ✓ | ✓ | ✓× | ✓ | ✓ | ✓ |
| May | ✓✓ | ✓ | ✓ | ✓ | ✓✓ | ✓ | ✓ | ✓ | ✓✓ | ✓ | × | ✓ | ×× | × | × | × |
| July | ×× | × | × | × | ✓✓ | ✓ | × | ✓ | ✓✓ | ✓ | ✓ | ✓ | ✓✓ | ✓ | ✓ | ✓ |
| August | ✓✓ | ✓ | ✓ | ✓ | ✓× | ✓ | ✓ | ✓ | ✓✓ | ✓ | ✓ | ✓ | ×× | × | × | × |
| October | ✓✓ | ✓ | ✓ | ✓ | ✓× | ✓ | ✓ | ✓ | ✓✓ | ✓ | ✓ | ✓ | ×× | × | × | × |
| December | ✓✓ | ✓ | ✓ | ✓ | ✓× | ✓ | ✓ | ✓ | ✓✓ | ✓ | ✓ | ✓ | ×× | × | × | × |

**Table S2** Results of the linear mixed-effects models prepared for the canopy cover of poplar, *Robinia* and pine forests. Pairwise comparisons were considered only if the full models explained a significant proportion of the variation of the data. Pairwise p-values were adjusted with the fdr method. *p<0.05.

| | March | | April | | May | | June | |
|---|---|---|---|---|---|---|---|---|
| | *F* | *p* | *F* | *p* | *F* | *p* | *F* | *p* |
| Forest type | 109.46 | <0.001* | 96.69 | <0.001* | 1.80 | 0.204 | 5.95 | 0.010* |
| | *t* | *p* | *t* | *p* | *t* | *p* | *t* | *p* |
| Poplar - *Robinia* | -5.17 | <0.001* | -12.72 | <0.001* | . | . | 2.79 | 0.018* |
| Poplar - Pine | 9.41 | <0.001* | -1.49 | 0.153 | . | . | -0.37 | 0.718 |
| *Robinia* - Pine | 14.59 | <0.001* | 11.22 | <0.001* | . | . | -3.15 | 0.018* |

| | August | | October | | December | | January | |
|---|---|---|---|---|---|---|---|---|
| | *F* | *p* | *F* | *p* | *F* | *p* | *F* | *p* |
| Forest type | 15.14 | <0.001* | 4.87 | 0.013* | 141.64 | <0.001 | 107.52 | <0.001* |
| | *t* | *p* | *t* | *p* | *t* | *p* | *t* | *p* |
| Poplar - *Robinia* | 3.05 | 0.005* | 0.51 | 0.608 | -5.21 | <0.001 | -2.60 | 0.014* |
| Poplar - Pine | -2.44 | 0.017* | -2.41 | 0.032* | 11.25 | <0.001 | 11.20 | <0.001 |
| *Robinia* - Pine | -5.49 | <0.001* | -2.92 | 0.018* | 16.46 | <0.001 | 13.80 | <0.001 |

**Table S3** Results of the linear models prepared for the effect of canopy cover on microclimatic variables in *Robinia* and poplar forests, and the one-sample t-tests on the effect of pine forests on microclimate as compared to adjacent open grasslands. *p<0.05.

| Robinia | *Estimate* | *t* | *p* | $R^2$ |
|---|---|---|---|---|
| DT TD | -0.14 | -4.17 | 0.001* | 0.52 |
| NT TD | 0.04 | 2.77 | 0.014* | 0.32 |
| DT HD | 0.35 | 3.45 | 0.003* | 0.43 |
| NT HD | 0.12 | 2.35 | 0.032* | 0.26 |

| Poplar | *Estimate* | *t* | *p* | $R^2$ |
|---|---|---|---|---|
| DT TD | -0.14 | -2.20 | 0.041* | 0.21 |
| NT TD | 0.01 | 0.26 | 0.795 | 0.01 |
| DT HD | 0.55 | 3.33 | 0.004* | 0.38 |
| NT HD | 0.23 | 2.18 | 0.042* | 0.21 |

| Pine | *Mean* | *t* | *p* |
|---|---|---|---|
| DT TD | -8.17 | -8.64 | 0.001* |
| NT TD | 1.38 | 3.53 | 0.003* |
| DT HD | 14.05 | 5.96 | <0.001* |
| NT HD | 0.79 | 0.39 | 0.703 |

**Table S4** Results of the linear mixed-effects models prepared for carbon and nitrogen contents of the soil of grasslands and poplar, Robinia and pine forests. Pairwise comparisons of habitat types were considered only within depth layers. Pairwise p-values were adjusted with the fdr method. *p<0.05.

| | Carbon content | | Nitrogen content | |
|---|---|---|---|---|
| | *F* | *p* | *F* | *p* |
| Habitat | 9.60 | <0.001* | 23.65 | <0.001* |
| Depth | 71.61 | <0.001* | 26.70 | <0.001* |
| 10–20 cm | *t* | *p* | *t* | *p* |
| Poplar - *Robinia* | -2.41 | 0.048* | 7.11 | <0.001* |
| Poplar - Pine | -3.52 | 0.004* | -1.19 | 0.324 |
| Poplar - Grassland | -2.95 | 0.018* | 0.74 | 0.506 |
| *Robinia* - Pine | -1.11 | 0.329 | -8.30 | <0.001* |
| *Robinia* - Grassland | -0.17 | 0.865 | -7.47 | <0.001* |
| Pine - Grassland | 1.11 | 0.329 | 2.11 | 0.072 |
| 70–80 cm | *t* | *p* | *t* | *p* |
| Poplar - *Robinia* | -2.37 | 0.048* | 3.09 | 0.008* |
| Poplar - Pine | -4.13 | 0.001* | -1.05 | 0.361 |
| Poplar - Grassland | -3.59 | 0.004* | -0.02 | 0.981 |
| *Robinia* - Pine | -1.76 | 0.151 | -4.14 | <0.001* |
| *Robinia* - Grassland | -0.85 | 0.436 | -3.59 | 0.002* |
| Pine - Grassland | 1.18 | 0.329 | 1.19 | 0.324 |

**Table S5** Results of the linear mixed-effects models prepared for the soil moisture content of grasslands and poplar, Robinia and pine forests over the course of a year. Centimetres from 20 till 120 denote depth layers. Pairwise p-values were adjusted with the fdr method. *p<0.05.

| | 20 cm | | 40 cm | | 60 cm | | 80 cm | | 100 cm | | 120 cm | |
|---|---|---|---|---|---|---|---|---|---|---|---|---|
| MARCH | F | p | F | p | F | p | F | p | F | p | F | p |
| Habitat type | 8.81 | <0.001 | 13.76 | <0.001 | 15.48 | <0.001 | 2.93 | 0.075 | 0.18 | 0.912 | 0.40 | 0.751 |
| | t | p | t | p | t | p | t | p | t | p | t | p |
| Grassland - Poplar | 4.53 | <0.001* | 2.77 | 0.011* | 0.46 | 0.903 | . | . | . | . | . | . |
| Grassland - *Robinia* | 3.66 | 0.003* | 1.57 | 0.144 | 0.10 | 0.923 | . | . | . | . | . | . |
| Grassland - Pine | 1.32 | 0.227 | -4.17 | <0.001* | -6.07 | <0.001* | . | . | . | . | . | . |
| Poplar - *Robinia* | -0.75 | 0.451 | -1.03 | 0.305 | -0.32 | 0.903 | . | . | . | . | . | . |
| Poplar - Pine | -2.78 | 0.014* | -6.01 | <0.001* | -5.65 | <0.001* | . | . | . | . | . | . |
| *Robinia* - Pine | -2.02 | 0.071 | -4.98 | <0.001* | -5.34 | <0.001* | . | . | . | . | . | . |
| APRIL | F | p | F | p | F | p | F | p | F | p | F | p |
| Habitat type | 27.37 | <0.001 | 10.43 | <0.001 | 32.78 | <0.001 | 85.26 | <0.001 | 18.86 | <0.001 | 13.99 | <0.001 |
| | t | p | t | p | t | p | t | p | t | p | t | p |
| Grassland - Poplar | 7.9 | <0.001* | 3.76 | <0.001* | 0.15 | 0.880 | -1.72 | 0.108 | -3.51 | 0.002* | -3.77 | <0.001* |
| Grassland - *Robinia* | 6.80 | <0.001* | 1.63 | 0.107 | -2.45 | 0.024* | -2.80 | 0.010* | -0.11 | 0.916 | 0.21 | 0.838 |
| Grassland - Pine | 3.31 | 0.002* | -2.41 | 0.027* | -9.30 | <0.001* | -12.90 | <0.001* | -6.89 | <0.001* | -5.49 | <0.001* |
| Poplar - *Robinia* | -0.96 | 0.340 | -1.84 | 0.083 | -2.25 | 0.033* | -0.94 | 0.352 | 2.95 | 0.006* | 3.44 | 0.002* |
| Poplar - Pine | -3.98 | <0.001* | -5.35 | <0.001* | -8.18 | <0.001* | -9.68 | <0.001* | -2.93 | 0.006* | -1.49 | 0.168 |
| *Robinia* - Pine | -3.02 | 0.004* | -3.50 | 0.002* | -5.93 | <0.001* | -8.75 | <0.001* | -5.87 | <0.001* | -4.93 | <0.001* |
| MAY | F | p | F | p | F | p | F | p | F | p | F | p |
| Habitat type | 10.63 | <0.001 | 33.85 | <0.001 | 50.95 | <0.001 | 52.92 | <0.001 | 27.06 | <0.001 | 31.91 | <0.001 |
| | t | p | t | p | t | p | t | p | t | p | t | p |
| Grassland - Poplar | 3.81 | <0.001* | -4.55 | <0.001* | -6.23 | <0.001* | -6.48 | <0.001* | -5.60 | <0.001* | -7.55 | <0.001* |
| Grassland - *Robinia* | 2.88 | 0.008* | -6.03 | <0.001* | -7.35 | <0.001* | -6.30 | <0.001* | -5.00 | <0.001* | -5.90 | <0.001* |
| Grassland - Pine | -1.70 | 0.111 | -9.60 | <0.001* | -11.70 | <0.001* | -12.21 | <0.001* | -8.33 | <0.001* | -8.01 | <0.001* |
| Poplar - *Robinia* | -0.81 | 0.422 | -1.28 | 0.206 | -0.97 | 0.336 | 0.15 | 0.883 | 0.53 | 0.599 | 1.43 | 0.188 |
| Poplar - Pine | -4.77 | <0.001* | -4.37 | <0.001* | -4.74 | <0.001* | -4.96 | <0.001* | -2.36 | 0.025* | -0.40 | 0.692 |
| *Robinia* - Pine | -3.97 | <0.001* | -3.09 | 0.003* | -3.77 | <0.001* | -5.11 | <0.001* | -2.89 | 0.008* | -1.83 | 0.107 |

| | 20 cm | | 40 cm | | 60 cm | | 80 cm | | 100 cm | | 120 cm | |
|---|---|---|---|---|---|---|---|---|---|---|---|---|
| JULY | F | p | F | p | F | p | F | p | F | p | F | p |
| Habitat type | 22.89 | <0.001 | 74.96 | <0.001 | 96.20 | <0.001 | 137.35 | <0.001 | 171.61 | <0.001 | 64.25 | <0.001 |
| | t | p | t | p | t | p | t | p | t | p | t | p |
| Grassland - Poplar | -0.78 | 0.436 | -10.92 | <0.001* | -12.23 | <0.001* | -15.51 | <0.001* | -17.99 | <0.001* | -10.82 | <0.001* |
| Grassland - *Robinia* | 5.23 | <0.001* | -2.06 | 0.051* | -7.59 | <0.001* | -11.29 | <0.001* | -13.66 | <0.001* | -8.88 | <0.001* |
| Grassland - Pine | -4.22 | <0.001* | -12.50 | <0.001* | -15.17 | <0.001* | -17.08 | <0.001* | -18.12 | <0.001* | -11.00 | <0.001* |
| Poplar - *Robinia* | 5.21 | <0.001* | 7.67 | <0.001* | 4.02 | <0.001* | 3.66 | 0.001* | 3.75 | <0.001* | 1.68 | 0.117 |
| Poplar - Pine | -2.98 | 0.005* | -1.37 | 0.174 | -2.54 | 0.013* | -1.36 | 0.179 | -0.11 | 0.910 | -0.15 | 0.879 |
| *Robinia* - Pine | -8.19 | <0.001* | -9.04 | <0.001* | -6.56 | <0.001* | -5.02 | <0.001* | -3.87 | <0.001* | -1.83 | 0.106 |
| AUGUST | F | p | F | p | F | p | F | p | F | p | F | p |
| Habitat type | 9.20 | <0.001 | 55.57 | <0.001 | 62.03 | <0.001 | 50.69 | <0.001 | 38.49 | <0.001 | 64.86 | <0.001 |
| | t | p | t | p | t | p | t | p | t | p | t | p |
| Grassland - Poplar | 1.35 | 0.180 | -9.10 | <0.001* | -9.57 | <0.001* | -9.13 | <0.001* | -7.95 | <0.001* | -10.47 | <0.001* |
| Grassland - *Robinia* | 3.97 | <0.001* | -3.83 | <0.001* | -3.83 | <0.001* | -7.84 | <0.001* | -7.64 | <0.001* | -9.78 | <0.001* |
| Grassland - Pine | -1.80 | 0.090 | -11.56 | <0.001* | -11.56 | <0.001* | -10.22 | <0.001* | -8.39 | <0.001* | -10.87 | <0.001* |
| Poplar - *Robinia* | 2.27 | 0.039* | 1.46 | <0.001* | 4.56 | 0.147 | 1.12 | 0.321 | 0.27 | 0.786 | 0.59 | 0.665 |
| Poplar - Pine | -2.74 | 0.016* | -2.14 | 0.036* | -2.14 | 0.048* | -0.94 | 0.351 | -0.38 | 0.786 | -0.35 | 0.728 |
| *Robinia* - Pine | -5.00 | <0.001* | -3.55 | <0.001* | -6.69 | 0.001* | -2.06 | 0.065 | -0.65 | 0.776 | -0.94 | 0.523 |
| OCTOBER | F | p | F | p | F | p | F | p | F | p | F | p |
| Habitat type | 16.90 | <0.001 | 72.25 | <0.001 | 50.59 | <0.001 | 18.67 | <0.001 | 16.42 | <0.001 | 16.26 | <0.001 |
| | t | p | t | p | t | p | t | p | t | p | t | p |
| Grassland - Poplar | 2.59 | 0.017* | -7.91 | <0.001* | -9.40 | <0.001* | -6.42 | <0.001* | -5.71 | <0.001* | -5.71 | <0.001* |
| Grassland - *Robinia* | 5.44 | <0.001* | 1.41 | 0.163 | -3.14 | 0.003* | -2.43 | 0.021* | -1.05 | 0.354 | -1.94 | 0.068 |
| Grassland - Pine | -2.12 | 0.038* | -12.26 | <0.001* | -10.41 | <0.001* | -5.71 | <0.001* | -5.32 | <0.001* | -5.58 | <0.001* |
| Poplar - *Robinia* | 2.47 | 0.019* | 8.07 | <0.001* | 5.42 | <0.001* | 3.46 | 0.002* | 4.03 | <0.001* | 3.27 | 0.003* |
| Poplar - Pine | -4.08 | <0.001* | -3.76 | <0.001* | -0.87 | 0.386 | 0.62 | 0.538 | 0.34 | 0.736 | 0.12 | 0.908 |
| *Robinia* - Pine | -6.55 | <0.001* | -11.84 | <0.001* | -6.29 | <0.001* | -2.84 | 0.009* | -3.70 | <0.001* | -3.15 | 0.003* |

| | 20 cm | | 40 cm | | 60 cm | | 80 cm | | 100 cm | | 120 cm | |
|---|---|---|---|---|---|---|---|---|---|---|---|---|
| DECEMBER | F | p | F | p | F | p | F | p | F | p | F | p |
| Habitat type | 9.98 | <0.001 | 4.56 | 0.006 | 9.05 | <0.001 | 36.13 | <0.001 | 25.78 | <0.001 | 30.96 | <0.001 |
| | t | p | t | p | t | p | t | p | t | p | t | p |
| Grassland - Poplar | 3.26 | 0.003* | 2.32 | 0.046* | 1.09 | 0.338 | 1.24 | 0.220 | -1.84 | 0.070 | -4.61 | <0.001* |
| Grassland - *Robinia* | 4.94 | <0.001* | 1.23 | 0.267 | -0.61 | 0.544 | -2.81 | 0.008* | -5.14 | <0.001* | -4.75 | <0.001* |
| Grassland - Pine | 3.52 | 0.002* | -1.70 | 0.140 | -4.44 | <0.001* | -9.30 | <0.001* | -8.26 | <0.001* | -9.41 | <0.001* |
| Poplar - *Robinia* | 1.46 | 0.225 | -0.95 | 0.348 | -1.47 | 0.220 | -3.50 | 0.001* | -2.86 | 0.008* | -0.13 | 0.900 |
| Poplar - Pine | 0.23 | 0.821 | -3.49 | 0.005* | -4.78 | <0.001* | -9.12 | <0.001* | -5.56 | <0.001* | -4.16 | <0.001* |
| *Robinia* - Pine | -1.23 | 0.268 | -2.54 | 0.040* | -3.31 | 0.003* | 5.62 | <0.001* | -2.70 | 0.010* | -4.03 | <0.001* |
| JANUARY | F | p | F | p | F | p | F | p | F | p | F | p |
| Habitat type | 3.51 | 0.020 | 19.33 | <0.001 | 9.80 | <0.001 | 8.81 | <0.001 | 1.59 | 0.200 | 3.72 | 0.015 |
| | t | p | t | p | t | p | t | p | t | p | t | p |
| Grassland - Poplar | 1.61 | 0.167 | 2.63 | 0.016* | -0.34 | 0.734 | 0.92 | 0.361 | . | . | -2.37 | 0.062 |
| Grassland - *Robinia* | -0.10 | 0.923 | 1.15 | 0.254 | -1.56 | 0.185 | -1.40 | 0.197 | . | . | -2.97 | 0.024* |
| Grassland - Pine | -2.12 | 0.113 | -5.58 | <0.001* | -5.21 | <0.001* | -4.43 | <0.001* | . | . | -1.75 | 0.170 |
| Poplar - *Robinia* | -1.48 | 0.172 | -1.28 | 0.246 | -1.05 | 0.355 | -2.01 | 0.072 | . | . | -0.52 | 0.605 |
| Poplar - Pine | -3.23 | 0.011* | -7.10 | <0.001* | -4.22 | <0.001* | -4.64 | <0.001* | . | . | 0.54 | 0.605 |
| *Robinia* - Pine | -1.75 | 0.167 | -5.83 | <0.001* | -3.16 | 0.005* | -2.62 | 0.021* | . | . | 1.06 | 0.441 |

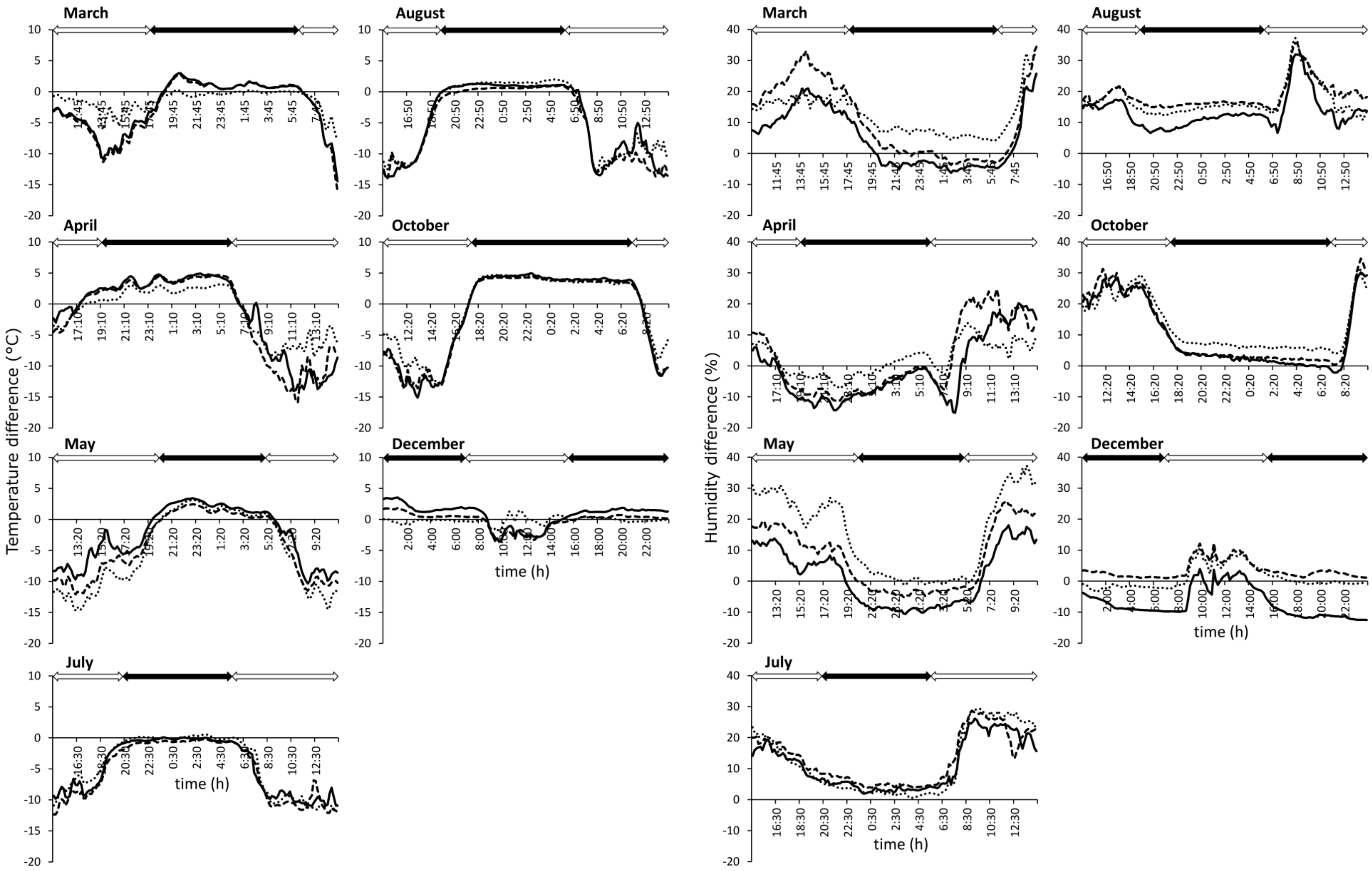


**Figure S1** Day-round microclimatic properties of poplar (dashed line), *Robinia* (dotted line) and pine forests (solid line) as compared to adjacent grasslands as base lines. Negative and positive values indicate that forests had respectively lower or higher values than grasslands at particular times of the day. Empty arrows: daytime (between sunrise and sunset), full arrows: night (between sunset and sunrise).

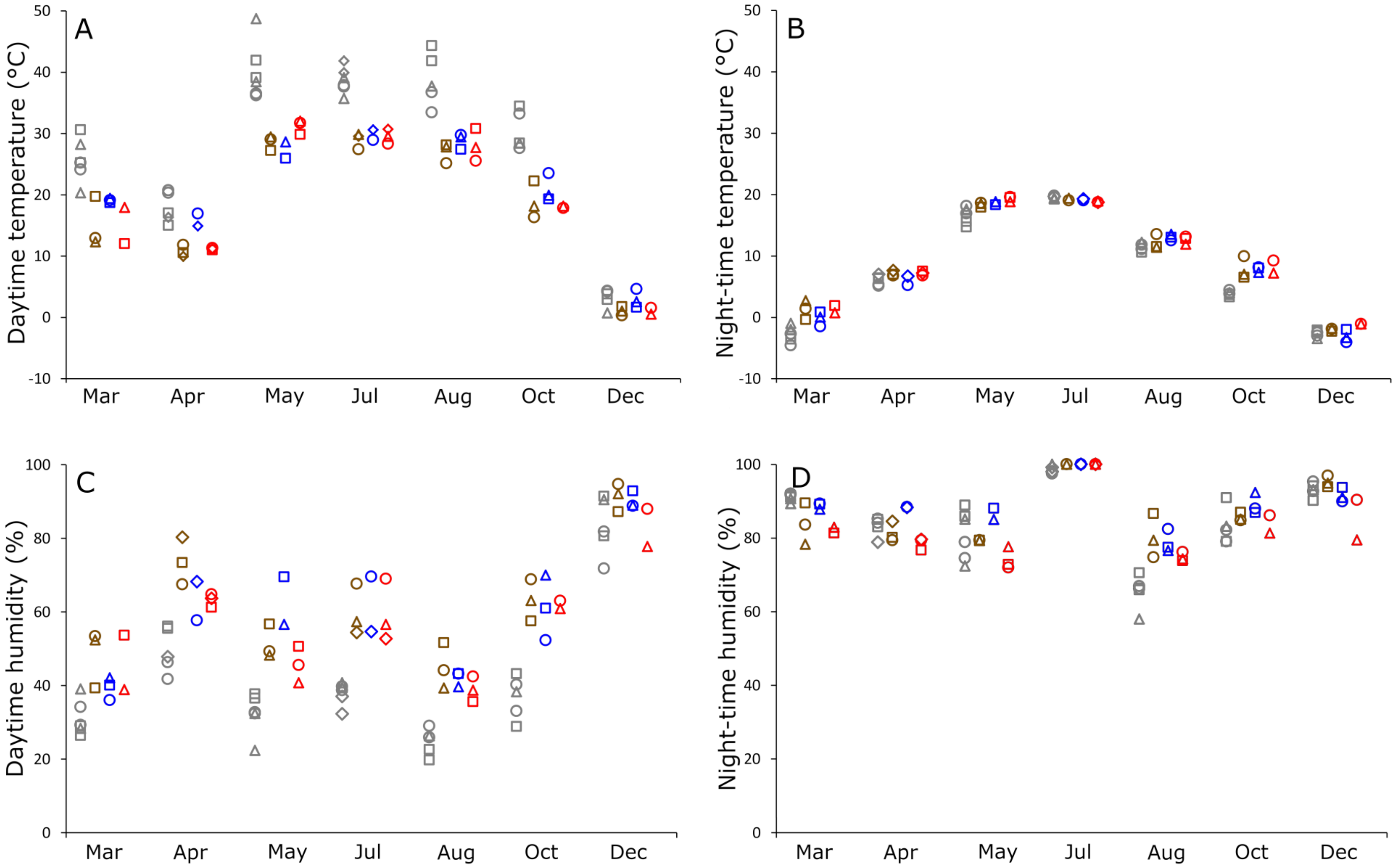


**Figure S2** Average non-standardized daytime (10 a.m. - 4 p.m.) and night-time (10 p.m. - 4 a.m.) temperature (A and B) and humidity (C and D) values of the sampling sites over the course of a year. Grey: grassland; brown: poplar forest; blue: Robinia forest; red: pine forest; circle: Méntelek; square: Fülöpháza; triangle: Ágasegyháza; diamond: Izsák.
